# Dynamic molecular epidemiology reveals lineage-associated single-nucleotide variants that alter RNA structure in Chikungunya virus

**DOI:** 10.1101/2021.01.17.427002

**Authors:** Thomas Spicher, Markus Delitz, Adriano de Bernardi Schneider, Michael T. Wolfinger

## Abstract

Chikungunya virus (CHIKV) is an emerging *Alphavirus* which causes millions of human infections every year. Outbreaks have been reported in Africa and Asia since the early 1950s, from three CHIKV lineages: West African, East Central South African, and Asian Urban. As new outbreaks occurred in the Americas, individual strains from the known lineages have evolved, creating new monophyletic groups that generated novel geographic-based lineages. Building on a recently updated phylogeny of CHIKV, we report here the availability of an interactive CHIKV phylodynamics dataset, which is based on more than 900 publicly available CHIKV genomes. We provide an interactive view of CHIKV molecular epidemiology built on Nextstrain, a web-based visualization framework for real-time tracking of pathogen evolution. CHIKV molecular epidemiology reveals single nucleotide variants that change the stability and fold of locally stable RNA structures. We propose alternative RNA structure formation in different CHIKV lineages by predicting more than a dozen RNA elements that are subject to perturbation of the structure ensemble upon variation of a single nucleotide.

## Introduction

Chikungunya virus (CHIKV) is an arthropod-borne *Alphavirus* of the family *Togaviridae* that causes millions of human infections every year, particularly in tropic and subtropic regions. CHIKV is the etiological agent of chikungunya fever, an acute febrile illness associated with joint pain, rash, and rarely neurological manifestations (1). CHIKV infection can culminate in chronic arthralgia and arthritis lasting up to several years. CHIKV cycles between vertebrate hosts and hematophagous arthropod vectors, predominantly *Aedes aegypti* and *Aedes albopictus* (2). In Africa CHIKV occurs in an enzootic, sylvatic cycle involving nonhuman primates as hosts, while in Asia CHIKV is mainly maintained in an urban cycle with direct human-mosquito-human transmission (3). The absence of a sylvatic life cycle in Asia suggests that CHIKV originated in Africa and was later carried to Asia (4).

### Geographical spread of Chikungunya virus

The first documented outbreak of CHIKV was in 1952 on the Makonde Plateau in the Southern Province of Tanganyika (present-day Tanzania) (5, 6). Since then, CHIKV has been emerging in Africa and Asia, with larger outbreaks in the 1960s and 1990s (7). In 2004, CHIKV re-emerged in Kenya and in 2005 an outbreak hit the island of La Réunion, which was the first time that CHIKV occurrences were reported in the southwestern Indian Ocean region (8). The virus, previously assumed to be non-fatal, caused several deaths and also affected neighboring islands including Mayotte, Madagascar, the Seychelles, Comoros, and Mauritius. Following the Indian Ocean islands outbreak, CHIKV spread independently into the Indian Subcontinent and Southeast Asia (9). In the Indian Ocean basin, CHIKV dissemination has been mediated by several mutations, the most prominent being A226V, an amino acid substitution that changes the protein structure of the membrane fusion glycoprotein E1 (10), resulting in increased transmission by *A. albopictus*. As *A. albopictus* is present in temperate regions, this adaptation also has implications on the geographical range of transmission, with CHIKV no longer being bound to tropical and subtropical latitudes (4). Reported cases of CHIKV in Italy, France, Mexico, and the USA, together with the expanding global distribution of *A. albopictus*, facilitated by climate change, raises public health concerns worldwide (11–13).

Early phylogenetic analyses suggested that CHIKV can be separated into three geographically disjoint lineages: West African (WA), East Central South African (ECSA) and Asian Urban Lineage (AUL) (14). Following the 2005 La Réunion outbreak, and the subsequent emergence of CHIKV in India, the existence of a fourth separate lineage, termed Indian Ocean Lineage (IOL), has been proposed (15). While the WA lineage is geographically isolated and shares deep ancestry with the other lineages, investigation of the relationships between taxa belonging to ECSA through phylogenetic inference revealed that this lineage in fact splits into three distinct geographically disjoint epidemic clades (16): The Middle African Lineage (MAL) gave rise to outbreaks in South America (17, 18) and Haiti (19) (South American Lineage, SAL), and the East African Lineage (EAL) gave rise to the IOL. The latter is predominantly found on the Asian continent, except for travel-related cases in which CHIKV has been imported to Europe and North America (3, 20, 21). The third ECSA-derived lineage includes the 1953 Tanganyika strain and encompasses a handful of isolates from Africa and Asia in a monophyletic group (Africa/Asia Lineage, AAL). The AUL, conversely, is considered a sister clade to all ECSA-derived lineages and has been circulating in Southeast Asia before spreading into Central America and many South American countries from 2013 (3, 22, 23). In 2013, the AUL lineage reached Brazil with the first autochthonous cases being observed in late 2014 (17). Around the same time span 2013/2014 it has emerged in the Caribbean, causing massive spread and nearly three million cases (24). Since then, AUL has spread to multiple regions in America. Interestingly, it is believed that CHIKV was present in the Americas in the 1800s, a courtesy of a spread through navigation, starting in the Caribbean and from there spreading into North and South America, although at the time CHIKV was mislabeled as another febrile disease, dengue (25).

### RNA structure conservation in Chikungunya virus genomes

CHIKV is a small, spherical, enveloped virus with a single-stranded, (+)-sense RNA genome of approximately 11.8 kb (2, 26) that contains a 5’ cap structure and a 3’ poly-A tail. The CHIKV genome contains two open reading frames (ORFs) that encode non-structural and structural proteins, respectively, as polyproteins that are post-translationally cleaved (27). The non-structural proteins (nsP1, nsP2, nsP3 and nsP4) constitute the viral replication machinery and are translated from the full-length genome, while the structural proteins (C, E1, E2, E3) form the virus particles and are produced from a subgenomic messenger RNA. The coding sequence is flanked by structured untranslated regions (UTRs) on both ends, which represent the most variable regions of the CHIKV genome (28). This divergence manifests in variable 3’UTR (and thus genome) lengths of individual CHIKV lineages, with ECSA-derived lineages being shorter than WA, and AUL comprising the longest isolates (15). It is plausible to propose patterns of coupled historic mutation and recombination events that eventually resulted in the plasticity observed in present day CHIKV isolates (29). Specific patterns and copy numbers of sequence-level direct repeats in the 3’UTR are characteristic of particular CHIKV lineages and likely represent adaptations of the virus to environmental constraints (30).

Like many other RNA viruses, CHIKV encodes not only viral proteins but also functional RNAs that mediate the viral life cycle. These structured RNAs are found in coding and noncoding regions of the viral genome. A specific fold is often a prerequisite for functional RNAs, and there are selective evolutionary pressures on maintaining these folds, both in coding and non-coding regions (31). As RNA structure is typically conserved at the level of secondary structures, structural homology can be interpreted as a result of evolutionary forces that act on particular RNAs, requiring them to maintain a critical set of structure-determining base pair interactions. In nature, this is achieved by compensatory mutations, i.e. those that maintain base-pair complementarity by a combination of two mutations, e.g. AU → GC, or consistent mutations that change only one pairing partner, e.g. AU → GU. While this kind of structural conservation of functional RNAs, which is also known a covariation, is an ubiquitous trait that is found in all domains of life, increased mutation rates render viruses particularly interesting in this context.

Although CHIKV is one of the best-studied viruses within the genus *Alphavirus*, knowledge of functional RNA elements and their specific association to viral pathogenesis and replication remains elusive. Unlike other RNA viruses that are characterized by structural conservation of a critical amount of functional RNAs in their UTRs, such as flaviviruses (32) or coronaviruses (33), genus-wide RNA structure conservation does not appear to be prevalent in alphaviruses. In this line, evidence for pervasive RNA structure conservation has not been observed among Sindbis virus (SINV), Venezuelan equine encephalitis virus (VEEV), and CHIKV (34), probably because the genomic location of recognized RNA structure motifs, such as packaging signals, are found at divergent locations in different alphaviruses. However, the apparent absence of covariation patterns among alphavirus species does not exclude the ability of individual species to form highly stable, functional structures. This has been recently affirmed in a genome-wide RNA structure probing study that could confirm known functional RNAs in CHIKV by SHAPE-MaP and characterize several highly structured, potentially functional RNA elements (35).

Likewise, the association between primary sequence and secondary structure in the terminal regions of CHIKV genomes has raised considerable research interest over the last years, mainly motivated by the observation that different CHIKV lineages maintain variable-length 3’UTRs that comprise specific patterns of sequence repeats. While earlier studies identified sequence repeat patterns in the 3’UTR of several alphaviruses (36), recombination by copy-choice mechanisms has been proposed to accelerate CHIKV adaptability, resulting in novel 3’UTR variants (30). In a recent study, we proposed an unambiguous association of sequence repeats in the 3’UTRs of different CHIKV lineages with evolutionarily conserved, structured and unstructured RNA elements (16). Current knowledge about the lack of functional conservation in alphaviruses suggests that potentially functional RNA elements evolved independently in each viral species.

An aspect related to the formation and specificity of RNA structure is the effect of single nucleotide variants (SNVs). These are mutations that can alter the RNA structural ensemble by mediating the base-pairing pattern, potentially resulting in an alternative fold and disrupted functionality. SNVs are sometimes associated with so called riboSNitches, i.e. RNA elements that are subject to perturbation of the structural ensemble resulting in large conformational changes (37, 38). Examples where such events can lead to disease phenotypes in human have been described in the literature (39–42).

### Molecular epidemiology reveals RNA structure-affecting SNVs

With the availability of large numbers of next generation sequencing data in public databases, multiple efforts to analyze and visualize the spread of infectious diseases have been made (43–45). Nextstrain (https://www.nextstrain.org) (46) is an open source project that makes pathogen phylogenetic data easily accessible to researchers and the interested public, thus facilitating research efforts in the field of pathogen evolution and epidemiology. Nextstrain allows to set up so called community builds that employ the Nextstrain software stack to construct custom real-time phylodynamics resources. Nextstrain community builds have become increasingly popular, and were used, for example, to highlight the genomic epidemiology of the 2018-2020 Ebola virus outbreak in DRC (47), and showcase the mutational dynamics of SARS-CoV-2 superspreading events in Austria (48). We report here the availability of a custom Nextstrain build for CHIKV that encompasses 924 publicly available genomes.

To better understand the evolutionary traits associated with functional RNA conservation among different lineages, we set out to use the CHIKV molecular epidemiology data for studying the impact of lineage-associated sequence variability on viral RNA structure. We were particularly interested in the structural divergence induced by fixed SNVs that are specific to particular lineages, and predict the existence of more than a dozen locally stable RNA elements in the coding regions of the CHIKV genome, whose structural ensemble is substantially altered by lineage-associated SNVs.

## Materials and Methods

### Taxon selection

We downloaded viral genome and annotation data from the public National Center for Biotechnology Information (NCBI) Genbank database (49) on 23 October 2020. We compiled all temporal and geographic metadata available in these genome records to create data sets for building the Nextstrain instance. Eight sequences of the data set were removed due to missing geographic location metadata or designation as a vaccine sequence. Metadata related to geographic location was associated with the United Nations geoscheme for consistency and labeled in the same way as in de Bernardi Schneider et al. (16). For the temporal analysis we identified two sequences with erroneous sampling dates in the Genbank record. Upon contacting the corresponding authors we were able to correct the dates of the NCBI accessions KX262991.1 and KY435454.1 to 2013 and 2014, respectively.

### Genetic distance

To calculate the genetic distance within and between Chikungunya lineages we estimated the evolutionary divergence over sequence pairs between groups as implemented in `MEGA X` under default settings (50, 51). The analyses were conducted using the Maximum Composite Likelihood model (52).

### CHIKV Nextstrain

We employed the workflow management system `Snakemake` (53) to build a pipeline for rapid deployment and reproducibility of the CHIKV Nextstrain build. In the first step of the `Snakemake` workflow, we retrieved metadata including collection date, country, and place of isolation (if available) from the Genbank records of all available CHIKV isolates. Each country was then assigned to one of the following regions: South East Asia, East Asia, South Asia, West Asia, Caribbean, Northern America, Central America, Southern America, Europe, Oceania, Eastern Africa, Middle Africa, Southern Africa, or Western Africa. In the next step, a file with lineage association for each isolate was created. Isolates with unknown lineage association were assigned to lineages via the the time-resolved phylogenetic tree produced by Nextstrain iteratively. The standard augur pipeline was then applied to construct all relevant data for Nextstrain visualization (46).

### RNA structure modulation via lineage-associated SNVs

For characterizing SNVs that affect CHIKV RNA structure formation we performed local RNA structure prediction with `RNALfold` (54) in the reference strain of our Nextstrain build (KT327163.2), limited to sequence lengths of 150 nt and filtered for thermodynamically stable structures. We required a free energy z-score of at least −2 when comparing to 1000 dinucleotide shuffled sequences of the same nucleotide composition, resulting in 138 locally stable candidate structures spread throughout the CHIKV genome. These were then intersected with Nextstrain genome diversity data, yielding a set of 759 candidate RNAs that overlap either one or more variable sites of the CHIKV genome. For each candidate RNA we computed the minimum free energy (MFE) of the non-mutated wild-type (WT) sequence as well as MFEs of all SNV mutants, assessed the basepair distance between WT and mutant MFE structures with `RNAdistance` from the `ViennaRNA` Package (55), employing a base-pair distance cutoff of 15, and filtered for variants that show (almost) complete fixation in one or more lineages. This yielded 14 locally stable RNA elements of the reference strain that overlap a total of 16 lineage-associated SNV, as listed in Table 4. Each SNV was then evaluated for its potential to alter the thermodynamic ensemble of RNA structures with the MutaRNA web server (56) using default parameters.

### Data availability

The CHIKV Nexstrain instance is available at https://nextstrain.org/community/ViennaRNA/CHIKV. The data can be downloaded by scrolling down to the bottom of the page and clicking on the “Download Data” link.

## Results

### Genetic distance between Chikungunya virus lineages

Nucleotide divergence analyses based on the average number of base substitutions per site can highlight the proximity among groups of taxa. To get an updated view of the distances within and between CHIKV lineages at the level of nucleotides, we computed the average evolutionary divergence over sequence pairs. Our results show that the nucleotide divergence within each individual lineage is relatively low (< 0.01) with an average of 0.006 (Table 1). The evolutionary divergence between lineages has an average value of 0.073 (Table 2). The WA lineage has a nucleotide divergence to the other lineages that ranges from 0.17-0.19, showing the highest divergence to all other lineages. Also, the major clades encompassing AUL and AUL-Am on one side, and EAL, IOL, MAL, and SAL on the other side present a nucleotide divergence ranging from 0.065-0.069 between each other. While the genetic divergence between these major clades confirms genotypes that have previously been described in the literature as well-defined lineages, i.e., WA, ECSA, and AUL, lower divergence can be observed between AUL-Am and AUL (0.011), as well as between SAL, MAL, EAL, IOL, AAL, sECSA, ranging from 0.006-0.031.

**Table 1.** Estimates of average evolutionary divergence over sequence pairs within Chikungunya virus lineages. The number of base substitutions per site from averaging over all sequence pairs within each group are shown.

| Lineage | Divergence |
| --- | --- |
| AUL-Am | 0.0012 |
| AUL | 0.0128 |
| SAL | 0.003 |
| MAL | 0.0107 |
| IOL | 0.0062 |
| EAL | 0.0011 |
| AAL | 0.0066 |
| WA | 0.0102 |

**Table 2.** Estimates of evolutionary divergence over sequence pairs between CHIKV lineages. The number of base substitutions per site from averaging over all sequence pairs between groups are shown. Analyses were conducted using the Maximum Composite Likelihood model (52) with 924 CHIKV nucleotide sequences encompassing the lineages/groups observed in this study. Lineages: Asian urban (AUL), AUL-America (AUL-Am), South America (SAL), Middle Africa (MAL), Indian Ocean (IOL), East Africa (EAL), Africa and Asia (AAL), Sister Taxa to ECSA (sECSA), West Africa (WA).

| Lineage | AUL-Am | AUL | SAL | MAL | IOL | EAL | AAL | sECSA |
| --- | --- | --- | --- | --- | --- | --- | --- | --- |
| AUL | 0.01108 |  |  |  |  |  |  |  |
| SAL | 0.06902 | 0.06622 |  |  |  |  |  |  |
| MAL | 0.06840 | 0.06548 | 0.02432 |  |  |  |  |  |
| IOL | 0.06988 | 0.06734 | 0.02954 | 0.02631 |  |  |  |  |
| EAL | 0.06763 | 0.06533 | 0.02574 | 0.02259 | 0.00584 |  |  |  |
| AAL | 0.06390 | 0.06067 | 0.03145 | 0.02887 | 0.03141 | 0.02807 |  |  |
| sECSA | 0.06302 | 0.06014 | 0.02957 | 0.02758 | 0.03038 | 0.02690 | 0.01918 |  |
| WA | 0.19383 | 0.19228 | 0.17696 | 0.17829 | 0.17917 | 0.17750 | 0.17443 | 0.17505 |

### A Nextstrain build for Chikungunya virus

In an attempt to provide a publicly available epidemiological and phylogeographical interactive visualization of CHIKV spread, we created a custom Nextstrain build that encompasses all currently available CHIKV genome data. Our community build is available via https://nextstrain.org/community/ViennaRNA/CHIKV and comprises 924 genomes, which represents a substantial increase compared to the 590 genomes considered in the previous most comprehensive study of CHIKV phylogeny (16). The Nextstrain phylogeny is based on a maximum-likelihood tree which is used to infer a timed tree with TreeTime (57) (Figure 1), making available the time of the most recent common ancestor (TMRCA) associated with each individual lineage/major clade of interest (Table 3).

**Fig. 1.**
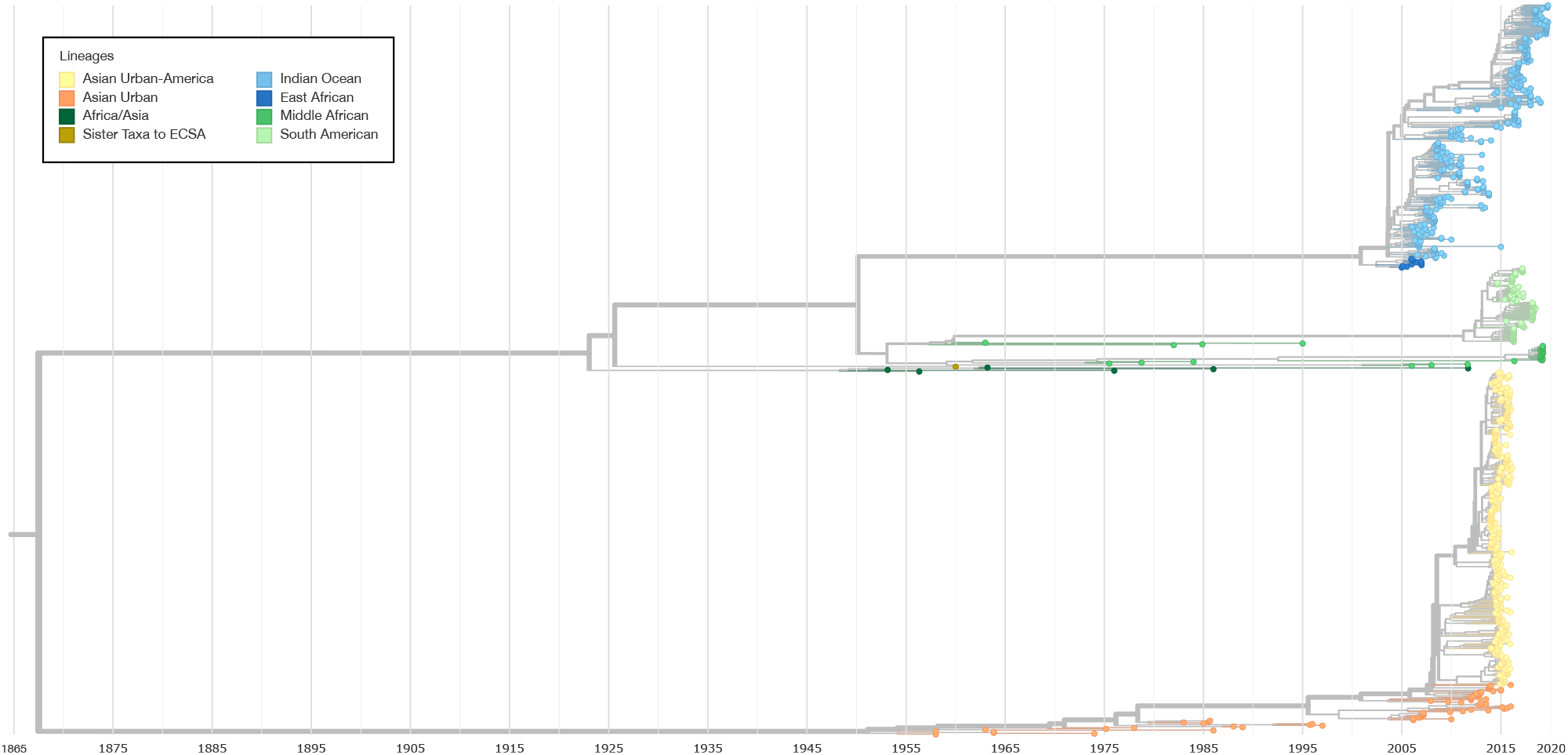
Nextstrain maximum likelihood phylogeny encompassing all CHIKV lineages except the West Africa lineage. The latter is located ancestral to all other clades with an inferred TMRCA date in 1645 (confidence interval 1625–1667, data not shown). Timing has been inferred by TreeTime.

**Table 3.** TMRCA estimates of CHIKV lineages, extracted from the Nextstrain instance. Dates are formatted as YYYY-MM-DD.

| Lineage | TMRCA | Date Confidence Interval | Year of First Isolation |
| --- | --- | --- | --- |
| AAL | 1948-04-17 | (1946-10-18,1949-12-14) | 1953 |
| AUL | 1951-02-05 | (1949-09-13,1953-01-04) | 1958 |
| AUL-Am | 2008-03-12 | (2007-10-24,2008-11-10) | 2013 |
| EAL | 2002-05-24 | (2001-02-15,2003-04-20) | 2005 |
| IOL | 2003-08-03 | (2002-10-20,2004-01-14) | 2006 |
| MAL | 1953-01-31 | (1951-05-20,1955-01-14) | 1962 |
| SAL | 2011-03-22 | (2009-06-24,2012-09-15) | 2014 |
| WA | 1954-01-16 | (1952-05-13,1955-09-26) | 1964 |

The CHIKV Nextstrain build allows the user to filter data and change the visualization according to preferences, utilizing a set of filters such as date ranges, multiple tree visualization layouts, feature filters and colors. Beside visualizing phylogeography-related characteristics, Nextstrain provides information about the diversity of the underlying sequence data as normalized Shannon entropy in a separate diversity panel. The analysis of nucleotide divergence within the full genome sequences available on the Nextstrain build enabled us to explore lineage-associated genomic variants that change the stability and fold of locally stable RNA structures *in silico*.

### Lineage-specific RNA structures

To better understand within-species RNA genotype-phenotype associations in viruses, wet set out to assess the impact of lineage-associated, evolutionary fixed SNVs on RNA structure formation in CHIKV. To this end, we performed local RNA secondary structure prediction in the reference strain of our Nextstrain build (KT327163.2, clustering with the AUL-Am lineage), aiming at characterizing structural elements that show increased thermodynamic stability, as expressed by z-score statistics. We intersected loci that fold into locally stable RNA structures with genome diversity data from Nextstrain to obtain regions of the CHIKV genome that both fold into stable RNA structures and contain one or more singlenucleotide mutations. Each SNV in this set was then characterized in terms of geographic spread and association to specific CHIKV lineages, as well as their base pair distance between non-mutated wild-type MFE structure and mutant MFE structure. Using base-pair distance to pre-filter variants that induce a substantial change in the global fold of the locally stable RNAs, we identified 12 candidate RNAs within the CHIKV coding regions that overlap one SNV, and two candidates that overlap two SNVs each. In total, we have 14 candidate RNAs and 16 fixed SNVs (Table 4).

**Table 4.**
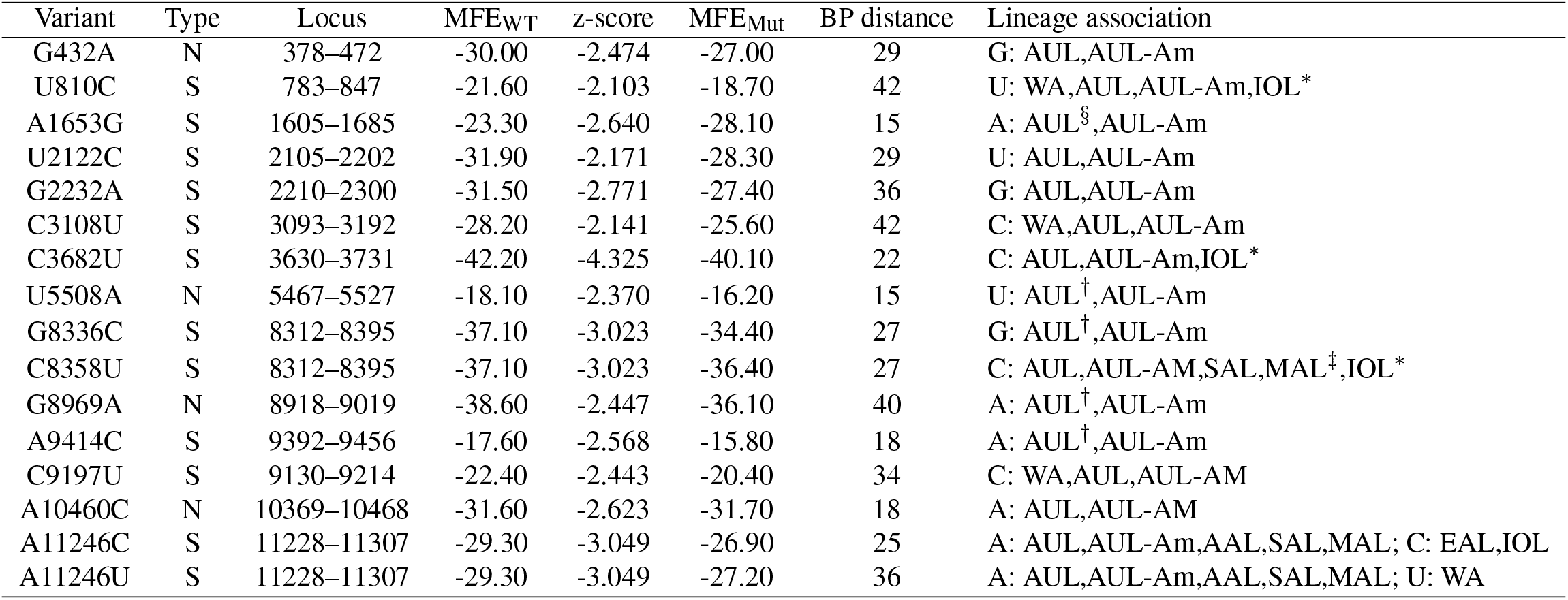
Lineage-specific single-nucleotide mutations that modulate the fold of locally stable RNA elements in the CHIKV genome, computed for the reference sequence of our Nextstrain build (accession KT327163.2). The Type column indicates whether a SNV induces a synonymous (S) or non-synonymous (N) mutation at the amino acid level. The Locus column lists global coordinates of the reference strain. Minimum fee energy (MFE) values for wild-type (WT) an mutant (Mut) are given in kacl/mol. Base-pair (BP) distance are computed between MFE structures of WT and Mut. *Subclade. ^§^Isolates collected from 1983 onwards. ^†^lisolates collected from 2006 onwards. ^‡^lisolates collected from 2019 onwards.

For each candidate RNA we quantified the effect of mutation-induced changes on the RNA structure ensemble with the `MutaRNA` webserver (56). In addition to comparing the characteristics of wild-type and mutant^1^ MFE structures, we assessed the impact of lineage-associated mutations on the entire thermodynamic ensemble of RNA structures by partition function folding.

We pick out two examples that exhibit interesting structural traits: The first example is a variant at nucleotide position 1653 (Figure 2), which has an Adenine (A, wild-type) in the majority of AUL and AUL-Am isolates, while a Guanine (G, mutant) is found at this position in all other lineages. The A1653G variant overlaps a locally stable RNA of 81 nt, whose wild-type sequence folds into a bulged stemloop structure with an MFE of −23.30 kcal/mol. The mutant folds into a three-way junction structure with an MFE of −28.10kcal/mol and an equilibrium frequency of 0.3 in the thermodynamic ensemble, suggesting that the mutant is thermodynamically more stable than the wild-type variant. The base pairing potential of wild-type and mutant sequences are depicted as a heat-map dot-plots as well as circular plots in Figure 2. These plots demonstrate the differences in base pairing patterns, highlighting that the SNV considerably modulates the fold of the RNA, thereby enabling the formation of alternatively stacked regions in the mutant that result in thermodynamic stabilization.

**Fig. 2.**
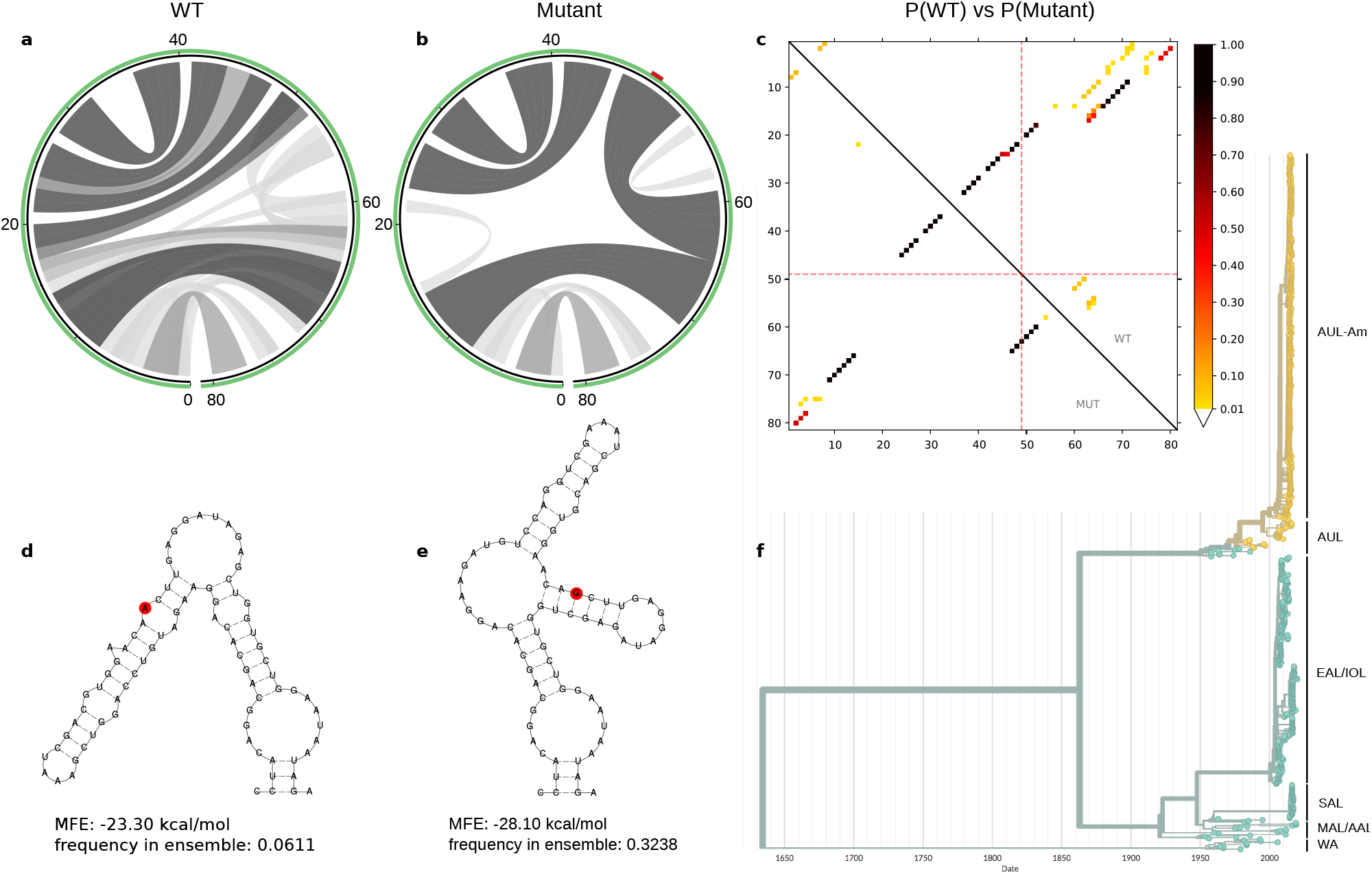
Ensemble properties of the 81 nt locally stable RNA element that contains the A1653G variant, represented as A49G here. Different patterns of base pair probabilities of the wild type (WT) and mutant are shown as circle plots in **a** and **b**, with darker arcs corresponding to increased base pairing probability. The SNV position in **b** is highlighted in red within the green circle that represents the RNA backbone. The dot plot in **c** shows the differences in base pairing potential of both WT (upper triangle) an mutant (lower triangle), with dark dots indicating high base pairing probability of particular sequence positions. Red dotted lines indicate the mutated position. Minimum free energy structures of the WT and mutant are shown in **d** and **e**, respectively, with the mutated position highlighted in red. The A1653G mutation results in a thermodynamic stabilization by 4.8kcal/mol. This mutation is specific to AUL, and in particular to AUL-Am isolates, highlighted in yellow in the time-resolved phylogenetic tree in **f**.

Another example encompasses multiple lineage-specific mutations at position 11246, as depicted in Figure 3. While the wild-type (comprising AUL, AUL-Am, AAL, MAL and SAL) has an A at this position, two other variants are clearly associated with different lineages: EAL and IOL have a Cytosine (C) and WA has an Uracil (U) at position 11246. While these variants result in a synonymous mutation at the amino acid level, they induce substantial changes for RNA structure formation. Position 11246 overlaps an RNA element of 80 nt with a wild-type MFE structure of − 29.30 kcal/mol and equilibrium frequency of 0.2156. The A11246C and A11246U variants are less stable, with MFE structures of −26.90 kcal/mol (equilibrium frequency 0.0650) and −27.70 kcal/mol (equilibrium frequency 0.1133), respectively. Intriguingly, all variants show varied base pairing patterns in the thermodynamic ensemble, where only 9 stacked base pairs of the closing stem are predicted with high probability.

**Fig. 3.**
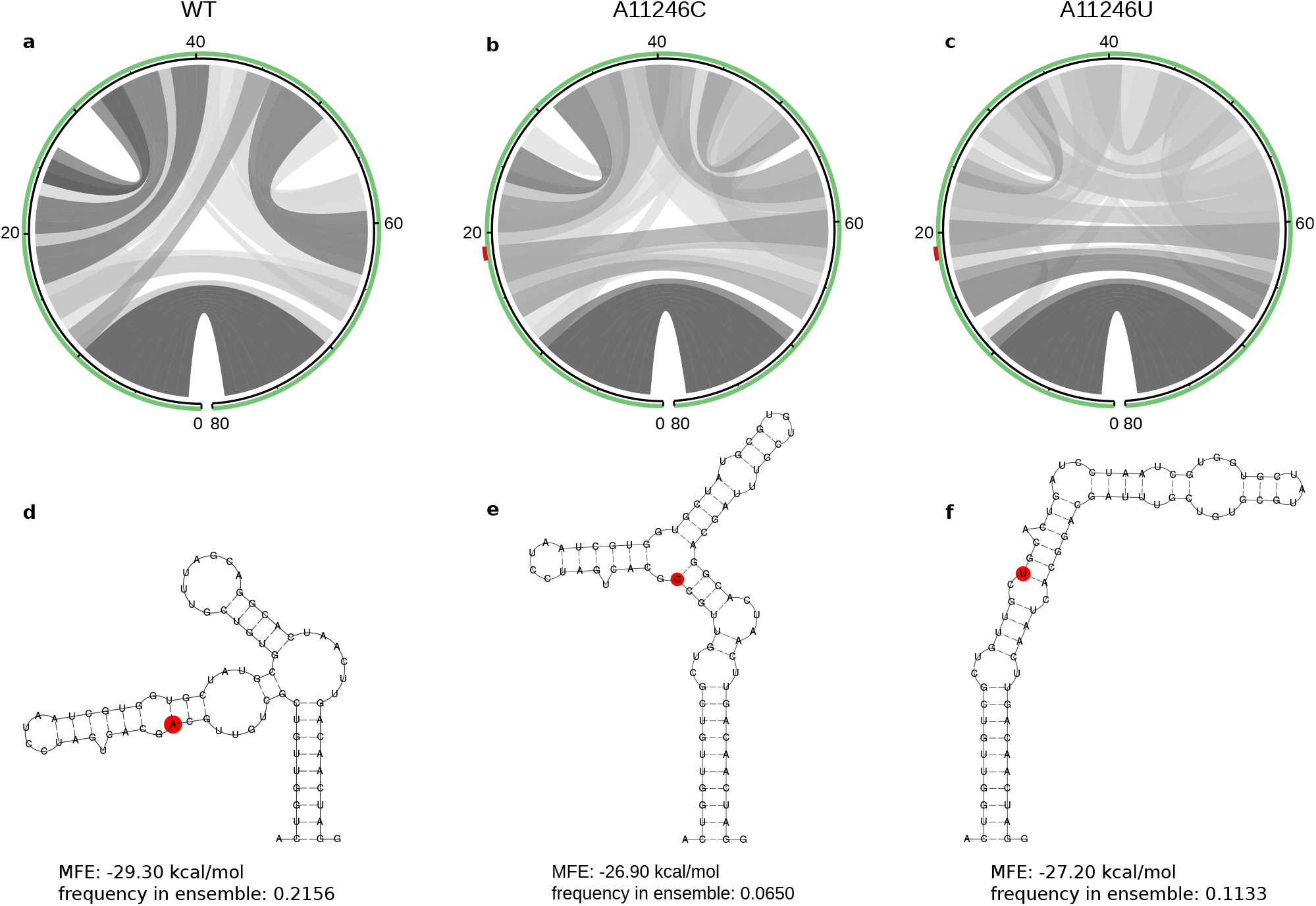
Aberration of the thermodynamic ensemble of a locally stable RNA element induced by mutations at position 11446, represented here as A19C and A19U. Structural diversity of the thermodynamic ensembles is shown in circle plots (**a-c**), where hue levels of gray represent pairing probabilities. The closing stem of 9 stacked base pairs is formed in all three variants, while the central parts of the RNA from positions 10–70 show varied base pairing patterns. The SNV at position 19 is highlighted in red on top of the green circles in **b** and **c** as well as in the minimum free energy structures **d-f**.

An interesting observation relates to a C-U mutation at position 10651, which is responsible for the E1-A226V mutation that has been associated with increased CHIKV transmissibility by *A. albopictus*. Although the C10651U mutation overlaps a locally stable region (positions 10594-10682 of the CHIKV genome, z=-3.73), the overall fold of this structure is not altered by the mutation. This suggests that the biological effect is mediated at the protein level rather than the RNA level.

## Discussion

In this contribution we address the question as to what can be learned about CHIKV genotype/phenotype associations by comparative approaches, bringing together different concepts of molecular epidemiology, phylogeny reconstruction, and computational RNA biology. To this end, we build on the Nextstrain (46) framework to provide an interactive phylodynamics resource of CHIKV that reveals spatiotemporal and epidemiological facets of global virus dissemination. Moreover, we employ established tools for RNA structure prediction based on the ViennaRNA (55) Package to infer lineagespecific structural traits.

While the nucleotide divergence within the observed CHIKV lineages is relatively low, this can be explained by the geographical constraint and the reduced collection period for novel lineages. Conversely, the divergence between lineages varies considerably. The highest divergence of the West African lineage when compared to the other CHIKV lineages can be explained by the observation that WA has been an isolated lineage that emerged decades before the more recent outbreaks (14).

However, the divergence between major clades and geographically delineated lineages does raise the question of whether the current nomenclature, as canonized by experts from the field when first identified, is not misguiding. IOL is an example of a well-established lineage in the literature that presents low divergence from its African origin, but it does introduce the A226V mutation in E1 which mediates the capacity of the virus to replicate in *A. albopictus* (58).

Most viral genotypes/subgenotypes/subclades are based on whole-genome nucleotide divergence of a specific percentage, usually determined by phylogeny showing clades/lineages of a defined ‘high’ statistical support (>70% bootstrap) (59, 60). Hepatitis B viruses, for example, are classified into genotypes and subgenotypes based on their monophyly, amino acid signatures, and genetic distance (61). In HIV, major distinct clades are classified as major genetic groups, with multiple subtypes within the genetic groups (62).

In the most recent comprehensive phylogeny of CHIKV to date, de Bernardi Schneider et al. (16) analyzed the three major lineages of CHIKV, AUL, IOL, and ECSA. These lineages were broken down based on their monophyly and geographic predominance. Here, we can see that although these strains can still be classified into distinct groups, there should be definite layers, such as lineages/genotypes and sublineages/subgenotypes.

Although we were not aiming at discussing a reclassification of CHIKV into a system independent of geography, we find the urge to bring to attention that a new coherent system should replace the current taxa classification. Such a new system could assist drug and vaccine development researchers to target specific genotypes or subgenotypes.

In an epidemiological context, the current lineage system, or, respectively, the way the strains are currently grouped, allows the inspection of major outbreak instances and looking at TMRCAs as a way to gauging when a lineage has been introduced in a region, causing outbreaks. From this perspective, our Nextstrain instance provides a reasonable amount of data to investigate deeper the outbreaks that have recently occurred. Importantly, the calculated TMRCA of the major clades is in accordance with previous studies (15). While before December 2013, local CHIKV transmission had not been identified in the Americas (24), our results suggest that the time to the most recent common ancestor for sequences in SAL and AUL-Am were 2011 and 2008, respectively. This result emphasizes the importance of increased surveillance, to identify the virus at the time of introduction, rather than at the time of pandemic (63). The consistency of the dates between Nextstrain TimeTree calculations and previously described TMRCAs in the literature is also reassuring of our ability to provide this additional information on the CHIKV Nexstrain instance.

Molecular epidemiology provides a detailed picture of the geographical spread and fixation of RNA variants in viruses and can be used in combination with *in silico* RNA structure prediction to study the structural divergence of different lineages. Owing to the error-prone replication machinery inherent to many RNA viruses, SNVs are created continuously and represent the constitutive driving force behind viral quasispecies (64). Although most of these mutations are considered neutral in an evolutionary context (65), a detailed understanding of the impact and functional associations of lineage-specific SNVs in viruses remain elusive. Single nucleotide mutations can lead to non-synonymous mutations at the amino acid level that result in potentially different protein function. Likewise, nucleotide mutations that culminate in synonymous mutations still have the capacity to alter RNA structure formation, leading to RNA phenotypes that can influence e.g. co-translational protein folding efficiency and thereby mediate viral gene expression patterns (66).

We asked whether Nextstrain can be utilized to infer novel insight into RNA structure-association of individual clades. As Nextstrain is particularly convenient for discriminating characteristics of viral clades/lineages, we set out to expand the sequence-centric approach to computation of lineage-associated structural traits. By combining Nextstrain genome diversity data with RNA structure prediction methods, we could associate sequence variants observed in different CHIKV lineages with alternative RNA structure formation. While we cannot associate lineage-specific SNVs with particular biological functionality at this point, the fixation of nucleotide mutations in certain CHIKV lineages suggests at least that these variants do not have detrimental influence on the viral fitness.

In summary, we show here that the combination of molecular epidemiology data with RNA structure prediction can help to gain insight into hitherto unresolved aspects of genotypephenotype associations within viral species. On a broader scale, specifically when applied to different viruses, this can augment our understanding of RNA structure evolution.

## Footnotes

1 Due to the selection of our reference strain, ‘wild-type’ refers to isolate KT327163.2, whereas ‘mutant’ refers to the respective SNV in all candidates listed in Table 4.

